# Preschool musicality is associated with school-age communication abilities through genes related to rhythmicity

**DOI:** 10.1101/2024.09.09.611603

**Authors:** Lucía de Hoyos, Ellen Verhoef, Aysu Okbay, Janne R Vermeulen, Celeste Figaroa, Miriam Lense, Simon E Fisher, Reyna L Gordon, Beate St Pourcain

## Abstract

Early-life musical engagement is an understudied but developmentally important and heritable precursor of later (social) communication and language abilities. This study aims to uncover the aetiological mechanisms linking musical to communication abilities. We derived polygenic scores (PGS) for self-reported beat synchronisation abilities (PGS_rhythmicity_) in children (N≤6,737) from the Avon Longitudinal Study of Parents and Children and tested their association with preschool musical (0.5-5 years) and school-age (social) communication and cognition-related abilities (9-12 years). We further assessed whether relationships between preschool musicality and school-age communication are shared through PGS_rhythmicity_, using structural equation modelling techniques. PGS_rhythmicity_ were associated with preschool musicality (*Nagelkerke*-R_2_=0.70-0.79%), and school-age communication and cognition-related abilities (R^2^=0.08-0.41%), but not social communication. We identified links between preschool musicality and school-age speech-and syntax-related communication abilities as captured by known genetic influences underlying rhythmicity (shared effect β=0.0065(SE=0.0021), *p*=0.0016), above and beyond general cognition, strengthening support for early music intervention programmes.

## Introduction

Human communication is a complex ability that enables us to develop and use language and cooperate with others, shaping communities and culture^1^. Communication with others requires speech-related (e.g. intelligibility and fluency of speech) and language-related (e.g. complexity of spoken grammar) skills, in addition to sufficient audible processing, while impairments thereof may result in communication disorders^2^. However, communication within a social setting, known as social communication or pragmatic language, also involves social interaction-related skills^3^. In particular, social communication requires (i) adequate language use, (ii) the adaptation of language to the listener or situation, and (iii) the adherence to conversational conventions (e.g. turn-taking or conversational rapport)^3^. Difficulties to communicate efficiently are, therefore, also characteristic of many mental health problems, including neurodevelopmental conditions^4^.

The acquisition of communication skills in typically developing children is influenced by social, linguistic, and cognitive factors^5,6^, but also other facets beyond traditionally defined general cognition and verbal skills. An understudied but developmentally important precursor of later (social) communication and language abilities is early-life musical engagement. Music is universal to human cultures^7^. Musicality (i.e. the suite of abilities involved in perceiving, producing and engaging with music including rhythmic abilities)^8^ and communication share universal aspects including acoustic, motor and perceptual skills, suggesting a shared biological basis^9,10^.

Rhythmic abilities emerge early in life and already new-borns can detect the beat in music^7^. In infancy, rhythmicity promotes social communication between the caregivers and the child^11,12^. In preschool children, rhythmic abilities are also related to phonological segmentation and phonological awareness shaping language acquisition^13^. More generally, early-life music engagement has a beneficial impact on later-life communication aspects such as oral and written language^14,15^, but also cognition and social skills^11^. Hence, deepening our understanding of links between early-life musicality, including rhythmic skills, and later (social) communication abilities may also help us to better understand the aetiological mechanisms underlying communication difficulties in children.

Genetic factors partially explain individual differences in both communication and musicality. Twin studies of communication-related traits have reported high heritability estimates^16^. Heritability estimates for pragmatic language (i.e. the use of language in conversation and social situations) and communication skills range from 58% to 79%^16^. Musicality-related traits are heritable too, with twin studies reporting a heritability of 50% for rhythm discrimination^17^. Recently, a genome-wide association study (GWAS) of self-reported beat synchronisation (i.e. a yes/no response to “Can you clap in time with a musical beat?”) was conducted in a large, well-powered sample of 606,825 individuals^18^. Using genetic variants present in at least 1% of the general population (indexed by single-nucleotide polymorphisms, SNPs), this study reported a SNP-based heritability of 13-16%^18^. The availability of this large GWAS^18^ enabled, for the first time, the study of genetic overlap between rhythmicity and other traits, demonstrating associations with language^19,20^, mental health^21^ and other musicality measures^22,23^, mostly in adult populations.

Given phenotypic links between early musicality and speech-language skills^14,15^, we hypothesised that these relationships may extend to communication phenotypes and manifest in shared genetic links, especially during an early developmental window. Compared to phenotypic analyses, investigating genetic relationships may add specificity and robustness, as genetic approaches can adjust for confounding influences affecting many complex traits^24^. Adopting a genomic approach has, thus, important implications for theoretical frameworks predicting links between musicality and language^14,25^, and for unravelling opportunities of support for children with communication difficulties.

Here, studying a large longitudinal sample of 6,737 children from the Avon Longitudinal Study of Parents and Children (ALSPAC)^26,27^, we, first, investigate whether polygenic load for rhythmicity is associated with a series of parent-reported preschool musicality and school-age communication abilities in a population-based sample. Next, we model structural relationships across associated phenotypes using a multivariate analysis framework. Finally, we study whether phenotypic relationships between preschool musicality and school-age communication are captured by the polygenic load for rhythmicity.

## Methods

### Participants

ALSPAC is a population-based longitudinal pregnancy-ascertained birth cohort from the United Kingdom^26,27^ (estimated birth date 1991-1992, Supplementary Note 1). Ethical approval for the study was obtained from the ALSPAC Ethics and Law Committee and the Local Research Ethics Committees. Consent for biological samples has been collected following the Human Tissue Act (2004). Informed consent for the use of data collected via questionnaires and clinics was obtained from participants following recommendations of the ALSPAC Ethics and Law Committee at the time.

### Phenotype information

Parent-reported measures of communication, social communication, musicality and nursery rhymes were assessed in ALSPAC children. Across all measures, at least 6,737 individuals had genetic and phenotypic data available. The study website contains details of all the available data through a fully searchable data dictionary and variable search tool (http://www.bristol.ac.uk/alspac/researchers/our-data).

For this study, we define a developmental preschool window (0-5 years) before children enter key stages of school curricula in the UK (i.e. preschool and reception years), compared to a developmental school-age window (> 5 years), where children enter educational key stages (i.e. year 1 and above). Once children start schooling, many predictors of language-related phenotypes, such as phonological awareness, become reciprocally shaped by reading experience and may no longer reflect underlying aetiological mechanisms^28^.

### Phenotypic measures

#### Nursery rhymes (preschool)

Parents answered whether they play pat-a-cake (i.e. a nursery rhyme where parent and children clap their hands together following the rhythm of the rhymes) or other clapping games with their child, representing an important developmental milestone^29^. Children’s ability was assessed at 0.5 and 1.5 years of age using a 3-point Likert scale: “Yes does often”, “Has only done once or twice”, and “Has not done yet”. Categorical scores were created aligning with developmental milestones^29^. At 0.5 years, the child was considered musically engaged if they had done the activity at least once or twice (1: “Yes does often” / “Has only done once or twice”), and non-musically engaged otherwise (0: “Has not done yet”). At 1.5 years, the child was considered musically engaged if they did it often (1: “Yes does often”) and non-musically engaged otherwise (0: “Has only done once or twice”/”Has not done yet”), as typically developing children should be able to play nursery rhymes by one year^29^.

#### Musicality-related measures (preschool and school-age)

Parent reports of their child’s ability to sing at least three songs, hum a tune, and clap in time with a musical beat were available at 5, 6 and 7 years using ALSPAC-specific questions assessing children’s development, based on a 3-point Likert scale. Categorical scores were created for each measure representing whether the child was considered musically engaged (1: “Yes can do well”/ “Yes but not well”) or not (0: “Has not yet done”).

#### Social communication measures (school-age)

Social communication measures were assessed by the parents using the Social and Communication Disorders Checklist (SCDC)^30^ total score at ages 8, 11, 14 and 17 years. The SCDC is a 12-item questionnaire (3-point Likert scale; 1: “Not true”, 2: “Quite/sometimes true”, 3: “Very/often true”) to be completed by parents about their children’s social interaction and communication skills. The SCDC total score is a summation of the 12 items demonstrating high consistency (Cronbach’s Alpha=0.86) and high reliability (0.84-0.93)^30^.

#### Communication measures (school-age)

Parents reported on children’s communication skills at 10 years using seven subscales of the Children’s Communication Checklist (CCC)^31^. The CCC is a 70-item questionnaire (3-point Likert scale, 1: “Certainly true”, 2: “Somewhat true”, 3: “Not true”). The subscales include (A) intelligibility and fluency, (B) syntax, (C) inappropriate initiation, (D) coherence, (E) stereotyped conversation, (F) use of conversational context, and (G) conversational rapport. In addition, the pragmatic communication score is created as a summary score using subscales C to G. CCC subscales have a moderate/high consistency (0.62-0.83) and high reliability (0.74-0.87)^31^. Within this study, we refer to subscales C and E as appropriate initiation and non-stereotyped conversation, respectively, to align with the direction of the effect (i.e. lower scores, more communication difficulties).

#### Verbal cognition-related measures (school-age)

Due to recent findings of genetic overlap between language-related traits and musicality^19^, we included measures of children’s verbal abilities (verbal IQ) and phonological working memory (nonword repetition). Both scores were assessed at 9 years and measured with an abbreviated form of the Wechsler Intelligence Scale for Children (WISC-III)^32^ and the Nonword Repetition Test (NWRT)^33^, respectively (Supplementary Note 2).

### Genetic information

Genotyping was performed using the Illumina HumanHap550 quad chip. Standard genetic quality control checks at the SNP and individual level were carried out in PLINK^34^ (Supplementary Note 3). After quality control, our study comprised 8,226 unrelated children (51% males) of European genetic ancestry and 465,740 SNPs with high-quality direct genotyping data, which were imputed to the Haplotype Reference Consortium reference panel (version r1.1) using the Sanger Imputation Server (https://imputation.sanger.ac.uk).

### Polygenic scores

#### Approach

Polygenic scores (PGS) are a weighted sum of alleles associated with a trait of interest that are carried by an individual^35^. A “discovery” GWAS of the trait of interest (here rhythmicity) is used to extract the respective weights for each genetic variant. The “target” sample (here ALSPAC) is used to calculate, for each individual, their genetic propensity score for the trait of interest based on the genetic variants they carry, and to run, across all individuals, an association analysis between such score and a set of phenotypes.

#### Discovery sample

In this study, we used GWAS summary statistics on self-reported beat synchronisation adjusted for sex, age and genetic quality control measures (hereafter referred to as rhythmicity)^18^ from 23andMe Inc., including 606,825 individuals of European descent (91,5% controls, 64% females). Note that the original authors validated this self-reported rhythmicity measure using PGS and an independent experiment^18^, and, additionally, this measure showed overlap with well-validated music tests in follow-up studies^23^.

#### Target sample

To generate PGS_rhythmicity_ in ALSPAC children, we extracted a set of common (minor allele frequency>1%) and well-imputed (imputation INFO score>0.8) HapMap 3 SNPs (N=985,350).

#### PGS calculation

PGS_rhythmicity_ were computed using PRS-CS^36^, a method that applies a continuous-shrinkage parameter to adjust the effect sizes of the genetic markers. Using these re-estimated effect sizes, PGSs were generated with PLINK^34^ and, subsequently, Z-standardised. Note that PRS-CS does not require the selection of a *p*-value, as required for clumping and thresholding methods^36^.

#### Association analysis

Logistic (binary phenotypes) or linear (continuous phenotypes) regression models were used to test for the association of ALSPAC phenotypes with PGS_rhythmicity_. Regression analyses were corrected for age, sex and the first ten ancestry-informative principal components. The variation explained by the PGS_rhythmicity_ was expressed in terms of regression R^2^ for continuous traits and, in analogy, *Nagelkerke-*R^2^ for binary traits (Supplementary Note 4).

#### Multiple-testing threshold

We computed the effective number of phenotypes across the initial set of 25 phenotypes. We carried out matrix Spectral Decomposition^37^, which identifies the number of independent phenotypes based on phenotypic correlations. This yielded a multiple-testing threshold of 2.5x10^-3^ (0.05/20 estimated independent phenotypes).

### Identification of phenotypic factor structures

We model phenotypic relationships for traits associated with PGS_rhythmicity_ (i.e. measures with PGS_rhythmicity_ *p*<0.05) using a data-driven approach (Supplementary Note 5). To do so, first, we estimated the number of factors using principal component analysis based on the phenotypic correlation matrix. Second, we split the sample into two random halves, matched for sex and missingness patterns. Third, we fitted an exploratory factor analysis (EFA) to the first half of the sample (N=3,048). To approximate the EFA factor structure, we retained standardised EFA factor loadings (λ), capturing at least 1% of the phenotypic variation (|λ|>0.1). Fourth, the identified EFA structure was used to inform subsequent confirmatory factor analysis (CFA), fitted in the other half of the sample (N=3,053). CFA model fit was assessed using the comparative fit index (CFI), the Tucker–Lewis index (TLI), the Root Mean Square Error of Approximation (RMSEA) and the Standardised Root Mean Square Residual (SRMR) parameters (Supplementary Note 5). EFA and CFA models were fitted using both orthogonal (varimax) and oblique (oblimin) rotation in *lavaan* (R::lavaan,v0.6-14)^38^.

To control for covariate effects, EFA and CFA were conducted with transformed phenotypes, adjusting for age, sex and the first ten ancestry-informative principal components (Supplementary Note 5). Note that joint analyses of multiple phenotypes within a factor analysis framework do not require adjusting for multiple-testing.

### Genetic characterisation of phenotypic structures

We tested whether the correlation between phenotypic factors was attributable to shared genetic variation with PGS_rhythmicity_ while keeping the identified CFA factor structure otherwise fixed. Specifically, we incorporated PGS_rhythmicity_ into the CFA model structure by applying a framework analogous to mediation analysis (Supplementary Figure 1)^39^. Note that the indirect effect within a mediation framework will estimate the shared genetic effect between two phenotypic factors as captured by PGS_rhythmicity_ (hereafter referred to as shared effect).

### Adjustment for genetic confounding

We generated rhythmicity GWAS^18^ summary statistics excluding genetic effects shared with the EA GWAS summary statistics^40^ (rhythmicity-EA) by using a GWAS-by-subtraction framework^41^. GWAS-by-subtraction is an approach that fits a Cholesky model in genomic structural equation modelling^42^ (R::genomicSEM, v0.0.5) based on GWAS summary statistics from two traits, in our case, rhythmicity and EA (Supplementary Figure 2). Within the EA GWAS, EA was measured in years of education adjusted for sex, year of birth, their interaction and genetic quality control measures^40^. To avoid sample overlap, ALSPAC and 23andMe individuals were excluded from EA summary statistics (personal communication with A. Okbay)^40^. Using the derived GWAS_rhythmicity-EA_, we created PGS_rhythmicity-EA_. Following guidelines by the original authors^41^, the effective sample size of the GWAS_rhythmicity-EA_ was estimated as N=143,800.

## Results

### Study design

To understand the relationships between preschool musicality and school-age (social) communication abilities, we studied up to 6,737 unrelated children of European descent from the ALSPAC cohort adopting a two-stage research design (Figure 1). Within the first stage, we identify measures that genetically overlap with PGS_rhythmicity_, including validation of ALSPAC-specific musicality measures. Within the second stage, we model the phenotypic structure across PGS_rhythmicity_-associated measures (PGS_rhythmicity_ *p*<0.05), focussing on relationships between preschool musicality and school-age communication as captured by the shared polygenetic load with PGS_rhythmicity_.

**Figure 1.**
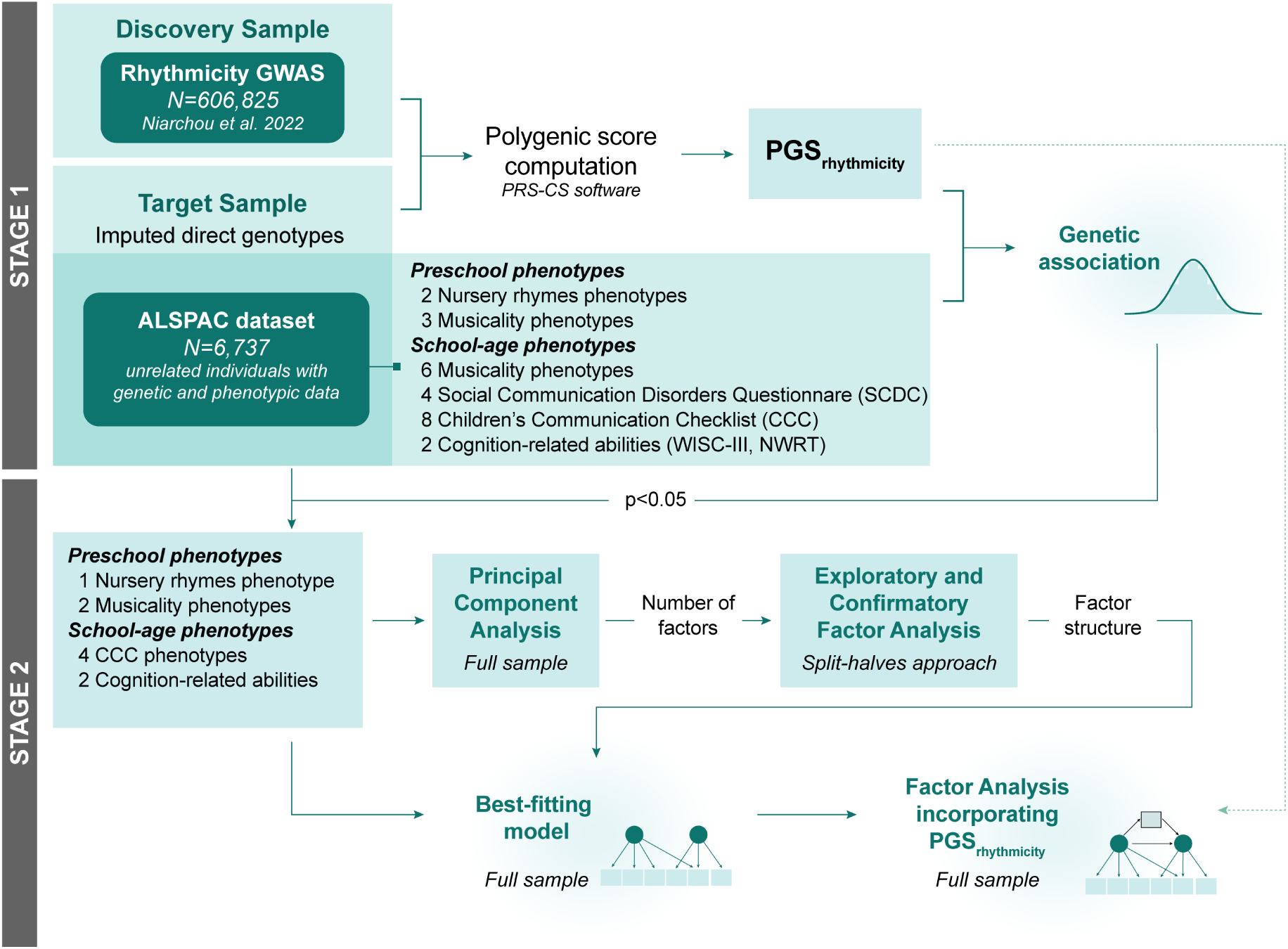
Study design. Analyses were carried out in 6,737 unrelated children of European descent from the Avon Longitudinal Study of Parents and Children (ALSPAC) cohort adopting a two-stage research design. In Stage 1, we constructed polygenic scores (PGS) for self-reported beat synchronisation^18^ (PGS_rhythmicity_) on ALSPAC individuals. Within Stage 2, we explored the phenotypic structure across phenotypes with PGS_rhythmicity_ p<0.05. To do so, we computed principal component analysis (PCA) and exploratory and confirmatory factor analysis, as described in the Methods. Finally, we mapped the PGS_rhythmicity_ to the factor structure, using methods analogous to mediation analysis.

Our analyses comprised a total of 25 measures (Table 1) assessed during preschool (6 months to 5 years, 5 measures) and school-age (6 to 17 years, 20 measures) years. Preschool measures include two infant ALSPAC-specific nursery rhyme measures and three early childhood ALSPAC-specific musicality measures. Six ALSPAC-specific school-age musicality measures were screened but showed ceiling effects (low cell counts) as the majority of children are already musically engaged at this developmental stage (Table 1). School-age communication measures comprised four SCDC social communication measures across childhood and adolescence, and eight mid-childhood CCC communication measures. Additionally, we studied school-age verbal cognition-related measures linked to rhythmicity, i.e. WISC-III verbal IQ and NWRT non-word repetition scores, capturing known links^19^.

**Table 1.**
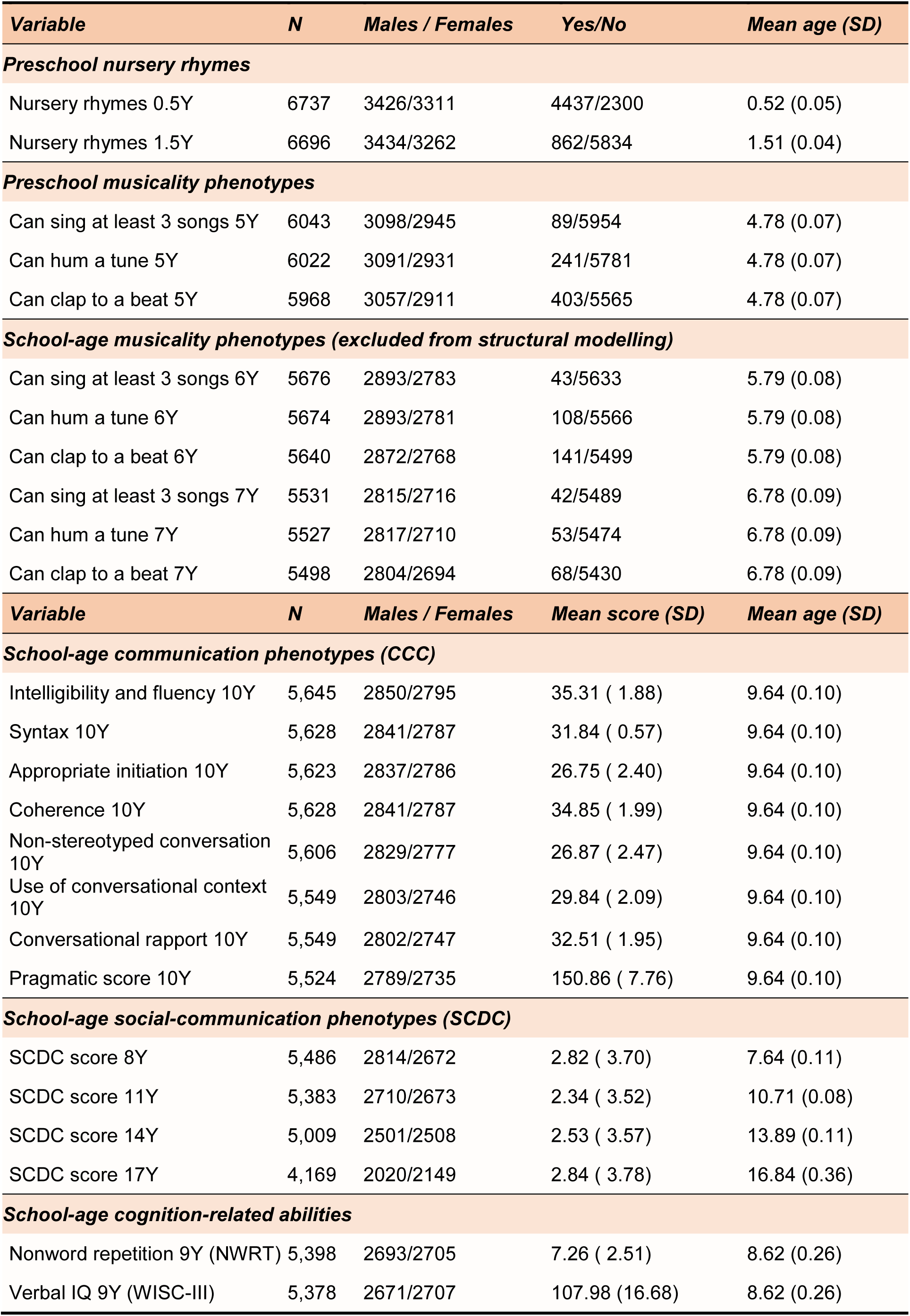
Descriptive information on the ALSPAC measures included in this study.

### Polygenic association analysis

During the first stage of our study, we screened the 25 phenotypes (Table 1) for polygenic overlap with rhythmicity (Figure 1). We found evidence for an association between PGS_rhythmicity_ and 11 measures at the nominal level (*p*<0.05, Figure 2, Supplementary Table 1), five of which passed the multiple-testing threshold (*p*<0.0025, Figure 2). Specifically, we observed positive links between PGS_rhythmicity_ and playing nursery rhymes at 6 months (β=0.097(SE=0.026),*p*=1.9x10^-4^) and between PGS_rhythmicity_ and preschool musicality-related abilities at 5 years, including humming a tune (β=0.23(SE=0.067),*p*=5.7x10^-4^) and clapping to a beat (β=0.22(SE=0.053),*p*=2.2x10^-5^). The identified genetic overlap with musicality-related measures confirms previous findings^18^ and extends known associations between PGS_rhythmicity_ and music engagement to an earlier developmental window. Genetic associations with PGS_rhythmicity_ were also observed for school-age communication abilities as measured by four CCC subscales at 10 years: intelligibility and fluency (β=0.12(SE=0.025),*p*=1.3x10^-6^), syntax (β=0.022(SE=0.0075),*p*=3.2x10^-3^), appropriate initiation (β=-0.068(SE=0.032),*p*=3.2x10^-2^) and conversational rapport (β=0.067(SE=0.026), *p*=9.3x10^-3^), although only the first one passed the multiple-testing threshold. The two school-age cognition-related measures, verbal IQ (β=- 0.55(SE=0.22),*p*=1.5x10^-2^) and nonword repetition (β=0.24(SE=0.034),*p*=5.1x10^-13^) also showed overlap with PGS_rhythmicity_, although only the latter passed the multiple-testing threshold. The overlap between rhythmicity and nonword repetition confirms, using individual-level genetic data, previously reported findings based on summary statistics^19^ (Figure 2, Supplementary Table 1). Overall, the proportion of variance explained by PGS_rhythmicity_ was modest (∼0.4-0.96%), yet in line with other studies of complex traits^43^. In contrast to speech-and language-related aspects of communication, there was little evidence for overlap between PGS_rhythmicity_ and social communication abilities (as measured with the SCDC) or the CCC subscales that are related to the use of language in a social context (Figure 2).

**Figure 2.**
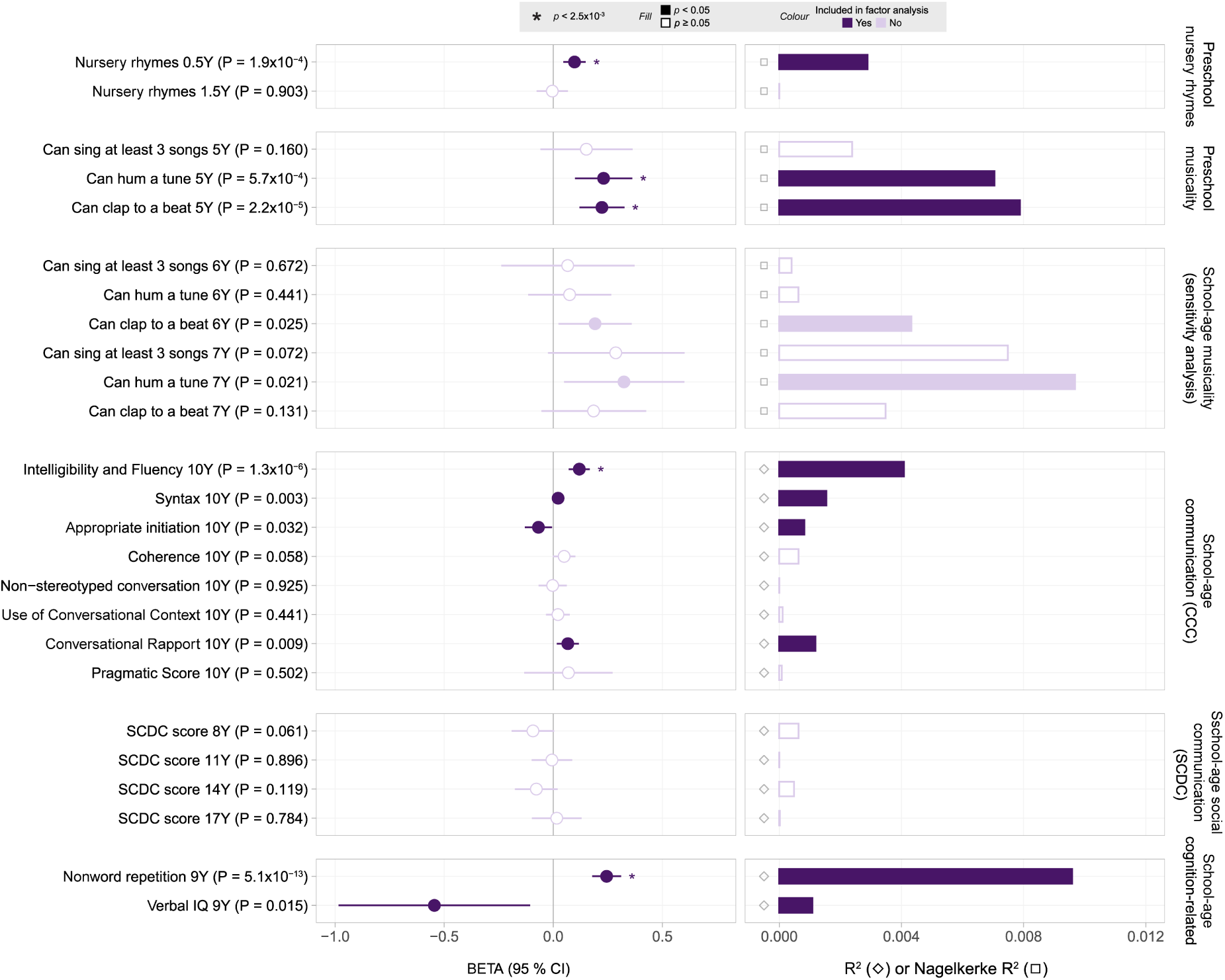
Effects of polygenic association with rhythmicity on ALSPAC measures. Beta estimates are shown as circles with their corresponding 95% confidence intervals. Variance explained for each phenotype is shown as bars and expressed as the regression R^2^ for continuous traits (represented by an empty grey diamond) and, in analogy, by *Nagelkerke*-R^2^ for binary traits (represented by an empty grey square). Filled circles/bars and empty circles/bars represent phenotypes with an association with PGS_rhythmicity_ of p<0.05 and p≥0.05, respectively. Estimates are shown in dark purple if they were included in subsequent factor analysis and in light purple otherwise. If a phenotype passed multiple-testing threshold of 2.5x10^-3^ this was indicated with an asterisk in the beta coefficient. A full table with the values for this analysis is available in Supplementary Table 1. Abbreviations: SCDC (Social Communication Difficulties Checklist), CCC (Children’s Communication Checklist).

Together, these findings suggest that genetic influences underlying rhythmicity are shared with preschool musicality and school-age cognition, working memory and speech-related communication abilities, consistent with overarching aetiological mechanisms.

### Identification and genetic characterisation of phenotypic structures

During the second stage of our study design, we studied the phenotypic structure across measures sharing a polygenic load with rhythmicity (PGS_rhythmicity_ *p*<0.05), conducting principal component and factor analyses (Figure 1). In particular, we aim to understand whether there is evidence for a developmental pathway linking preschool musicality to school-age communication through genes shared with rhythmicity. Note that we specifically selected preschool musicality measures, as school-age musicality measures showed ceiling effects and might be affected by reverse causation due to schooling^28^.

Firstly, we modelled the relationships across PGS_rhythmicity_-associated measures adopting a data-driven approach (Methods). We identified three phenotypic dimensions using principal component analysis, as reflected by three eigenvalues above one (Supplementary Figure 3). We, subsequently, fitted a three-factor EFA model with and without correlated factors and confirmed the identified structure in CFA using a split-halves approach (Figure 1). The identified three-factor CFA model, allowing for correlation between factors, showed a good model fit in the full sample (CFI=0.97, TLI=0.95, RMSEA=0.03, SRMR=0.02, Figure 3A). A preschool musicality factor (F1) described phenotypic variation of preschool ability to clap to a beat (λ=0.55(SE=0.041)) and hum a tune (λ=0.52(SE=0.039)) at 5 years. The school-age verbal cognition factor (F2) captured variation within nonword repetition (λ=0.61(SE=0.022)) and verbal IQ (λ=0.64(SE=0.022)) at 9 years. A school-age communication factor (F3) explained variation across three of the CCC subscales at 10 years: intelligibility and fluency (λ=0.57(SE=0.019)), syntax (λ=0.58(SE=0.019)) and conversational rapport (λ=0.39(SE=0.018)). Inter-factor correlations with F1 were modest (r_F1,F2_=0.21(SE=0.028), r_F1,F3_=0.27(SE=0.029)), while the correlation between F2 and F3 was strong (r_F2,F3_=0.57(SE=0.024)). Thus, PGS_rhythmicity_-associated measures are captured by three overarching phenotypic dimensions and, consequently, underlying association patterns might be shared across these dimensions, too.

**Figure 3.**
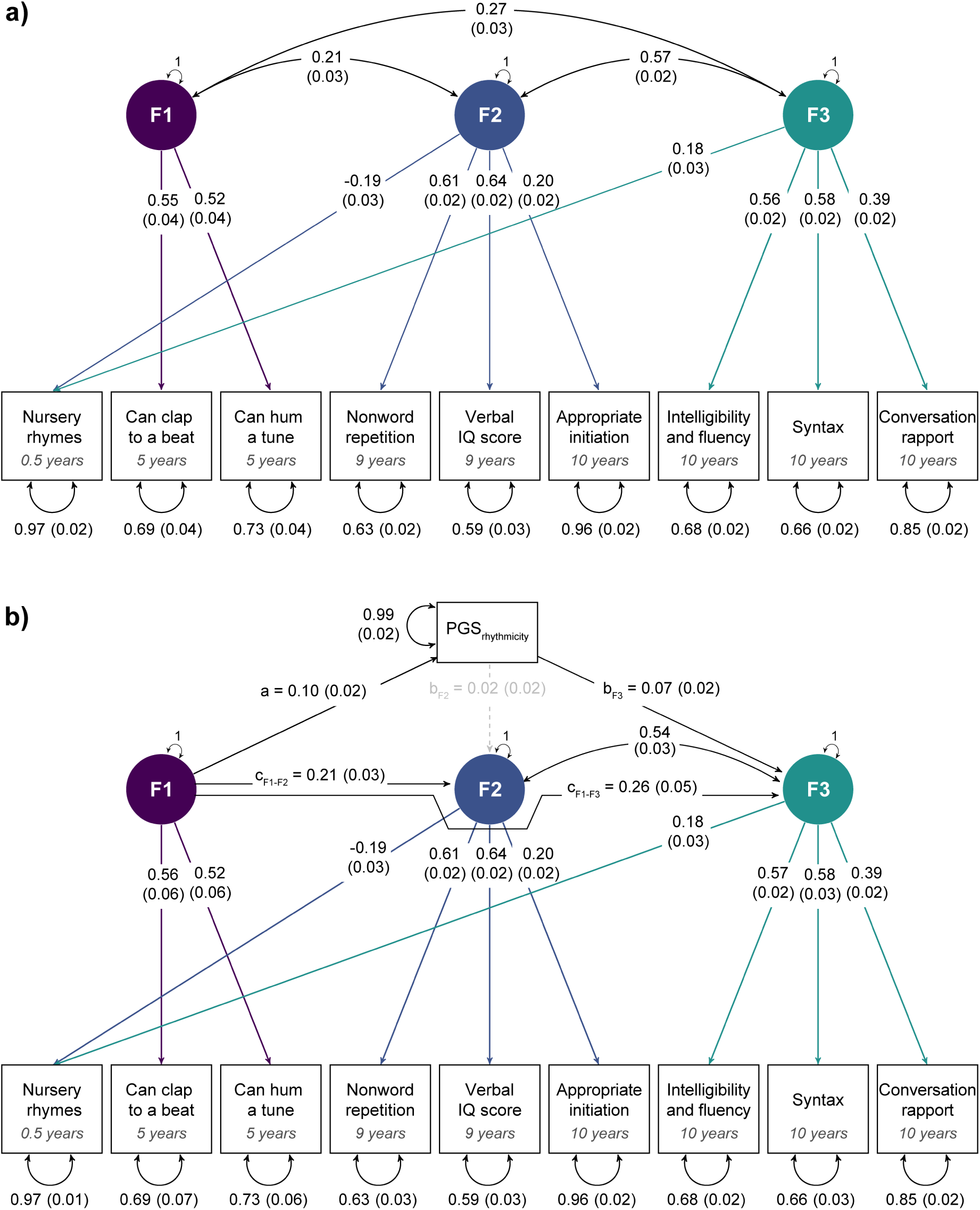
Phenotypic relationships between preschool musicality, school-age cognition-related skills and communication skills. **a)** Confirmatory factor model of phenotypes sharing genetic links with PGS_rhythmicity_. Estimates are shown with their corresponding SEs. Observed measures are represented by squares and latent variables by circles. Coloured single-headed arrows define factor loadings with p≤0.05. Double-headed black arrows represent the variance of each phenotype and factor correlations. The CFA model provided an optimal model fit (CFI=0.97, TLI=0.95, RMSEA=0.03, SRMR=0.02). **b)** Genetic characterisation of phenotypic relationships between F1 on F2 and F3 explained by shared genetic links with PGS_rhythmicity_. Estimates are shown with their corresponding SEs. Observed measures are represented by squares and latent variables by circles. Coloured single-headed arrows define factor loadings with p≤0.05. Double-headed black arrows represent the variance of each phenotype and factor correlations. Grey dotted and black solid single-headed arrows define relationships between factors and with PGS_rhythmicity_ with p>0.05 and p≤0.05, respectively. The shared genetic effect between F1 and F3, as captured by PGS_rhythmicity_, is estimated as a*b_F3_ and the total effect between F1 and F3 as a*b_F3_ + c_F1-F3_. The shared genetic effect between F1 and F2, as captured by PGS_rhythmicity_, is estimated as a* b_F2_ and the total effect as a* b_F2_ + c_F1-F2_.

Secondly, following the identification of phenotypic structure across these PGS_rhythmicity_-associated measures, we examined whether polygenic load for PGS_rhythmicity_ can capture overarching factor relationships, accounting for overlap with verbal cognition and phonological working memory (Supplementary Table 1, Figure 2). Adopting an analytical framework analogous to mediation analysis (Methods), we dissected the relationships between the three phenotypic factors (F1, F2 and F3) and modelled association effects shared with and without PGS_rhythmicity_. Specifically, we dissected the association of preschool musicality (F1) with both school-age verbal cognition (F2) and school-age communication (F3), as captured by polygenic influences underlying rhythmicity (Figure 3B). After adding PGS_rhythmicity_ to the CFA model structure, the model fitted the data similarly well (CFI=0.94, TLI=0.90, RMSEA=0.03, SRMR=0.02). Note that the factor structure of the model remained otherwise unchanged, such that estimates of the correlation between school-age verbal cognition (F2) and school-age communication (F3), as well as all other factor loadings, were virtually identical (Figure 3A vs Figure 3B).

Our analyses demonstrated that the association between preschool musicality skills (F1) and school-age communication skills (F3) (total effect: β=0.27(SE=0.047),*p*=1.59x10^-9^) was partially accounted for by PGS_rhythmicity_ (shared effect: β=0.0065(SE=0.0021),*p*=1.58x10^-3^, Supplementary Table 2). In contrast, there was little evidence that genetic variation shared with PGS_rhythmicity_ contributes to the association between preschool musicality (F1) and school-age verbal cognition (F2) (shared effect: β=0.0015 (SE=0.0022), *p*=0.49; total effect: β=0.21(SE=0.033), *p*=2.35x10^-11^). Within the structural model, all analyses were adjusted for each other. These association patterns show that preschool musicality is linked to the development of cognition-versus communication-related abilities, as captured by genetic load contributing to rhythmicity in large genome-wide studies^18^. As the proportion of explained phenotypic variation by PGS_rhythmicity_ for each phenotype, consistent with other studies^44^, was small (Figure 2), investigating the phenotypic structure across PGS_rhythmicity_-associated measures using structural models increases the power of our study.

### Adjustment for genetic confounding

Families with higher socioeconomic status (SES) may have easier access to music training, and there is evidence for an association between musicality and EA^45^, a proxy of SES with genetic contributions^40^. Consequently, we performed additional analyses to rule out genetic confounding by EA that may, potentially, affect links between preschool musicality and school-age social communication. We applied GWAS-by-subtraction techniques^41^ to create PGS_rhythmicity-EA_ (Methods, Supplementary Figure 2), thereby removing genetic variance from the rhythmicity GWAS shared with the EA, and repeated the PGS and structural equation modelling analyses as described above.

Association analysis using PGS_rhythmicity-EA_ (Stage 1, Figure 1) identified a similar pattern of genetic overlap across the studied measures, compared to analyses using PGS_rhythmicity_, especially for associations passing the multiple-testing threshold (Supplementary Figure 4, Supplementary Table 3). In addition, the phenotypic factor structures and association patterns remained robust (Supplementary Figure 5, Supplementary Table 4), suggesting that our findings are unlikely to reflect genetic effects shared with educational attainment, consistent with previous findings^18^.

Thus, genetic links between latent preschool musicality (F1) and school-age communication (F3), but not verbal cognition (F2), are partially attributable to genetic variation underlying rhythmicity(beat synchronisation)^18^, independent of genetic variation contributing to EA.

## Discussion

Investigating a UK population-based sample of unrelated children, this study showed that (i) preschool musicality and school-age communication abilities share genetic links with polygenic load for rhythmicity and that (ii) this overlap can partially explain phenotypic links between preschool musicality and school-age communication in population-based cohorts. These findings are in support of aetiological mechanisms, where preschool musical engagement, as captured by genes underlying rhythmicity, contributes to children’s school-age communication abilities during mid-childhood, above and beyond influences underlying general cognition.

The association of PGS_rhythmicity_ with early musicality measures spanning 6 months to 5 years (based on parent reports), including playing nursery rhymes at 6 months, clapping to a beat and humming a tune at 5 years, demonstrates that the genetic underpinnings of being able to clap to a beat in adulthood are transferable across the lifespan. Moreover, the association with PGS_rhythmicity_ validates these ALSPAC measures phenotypically, given the previous link of PGS_rhythmicity_ with objective tests of rhythm ability^23^.

The observed associations between PGS_rhythmicity_ and language-and communication-related phenotypes converge with previous studies reporting a phenotypic association between spoken language and rhythmic abilities in children^14^, extending them to communication abilities. Rhythm perception and production have been previously identified as predictors of phonological awareness, while melody perception has been linked to grammar acquisition^46^. Thus, our results are in line with two postulated frameworks the *Musical Abilities, Pleiotropy, Language, and Environment* (MAPLE) framework^14^ and the *Atypical Rhythm Risk Hypothesis*^25^, which discuss the potential role of rhythmicity in language-and communication-related traits.

The correlation between the factors of preschool musicality and school-age communication was partially accounted for by shared genetic effects with PGS_rhythmicity_. Consistently, impaired rhythmic skills and timing have been highlighted as an early predictor of atypical developmental cascades^47^ that may contribute to later language and communication difficulties. Given that phenotypic relationships are largely similar to genetic relationships^48^, our findings also strengthen the support for musical, and in particular rhythm, training during early childhood as an intervention to improve school-age speech-and syntactic language abilities^49,50^. This might be particularly relevant for children with fluency-related disorders, characterised by speech-flow interruptions^2^.

The factor for preschool musicality was also correlated with a school-age cognition-related factor. However, in contrast to school-age communication, there was little evidence for an association through shared genetic effects with PGS_rhythmicity_. Given comparable power (i.e. similar sample numbers) and cohort design, aetiological links between preschool musicality and school-age communication skills might be distinct from processes shaping working memory and cognitive development. However, larger samples are required to show a difference in effect, based on non-overlapping 95% confidence intervals.

This study has multiple strengths. First, we introduced a multivariate modelling framework combining PGS analyses with mediation and factor analytic techniques. This approach allows us to compare and combine PGS_rhythmicity_-related associations across a wide range of communication abilities. Hence, our findings based on structural models enhance the understanding of relationships across large phenotypic dimensions beyond individual PGS association analyses. Second, we characterise and validate the covariance structure between identified phenotypic factors by disentangling shared polygenic load with rhythmicity. In particular, this PGS-based study draws power from the large rhythmicity GWAS analysis^18^, increasing evidence for the specificity of the identified association between preschool musicality and school-age speech-related communication abilities. Still, the presented PGS association analyses need replication in an independent population-based cohort, which is challenging given the scarcity of longitudinal birth cohorts with genomic data and both (social) communication and musicality phenotypes, especially for individuals of non-European ancestry. Third, the longitudinal order phenotypes, split across preschool and school-age developmental windows, makes reverse causation unlikely. Fourth, genetic links between preschool musicality and school-age speech-related communication are robust to genetic confounding and independent of genetic effects related to EA. Thus, the link between preschool musicality and school-age speech-related communication is highly consistent with underlying causal mechanisms.

Nonetheless, our work has several limitations. First, ALSPAC, like other longitudinal cohorts, is plagued by loss to follow-up, and, consequently, findings might be affected by attrition bias. However, as presented findings are independent of genetic factors contributing to EA in the studied children, such bias is less likely. Second, association analyses of individual measures with PGS_rhythmicity_ were conducted using untransformed phenotypes as outcomes, while factor analytic approaches were carried out with transformed measures (adjusting different outcomes for different covariates). Consequently, we cannot exclude transformation bias, although findings were highly consistent across both stages. Third, the lack of association between PGS_rhythmicity_ and social-communication phenotypes may reflect a lack of power. However, speech-related and social communication phenotypes were both investigated with a comparable study design and, for CCC measures, also questionnaire design, rendering power an unlikely explanation. Instead, differences in genetic association profiles may pinpoint differences in aetiological mechanisms. Nonetheless, given the broad scope of human musicality^8^, there might be genetic links between social communication and musicality beyond rhythm production and perception (as captured by beat synchronisation). Likewise, other aspects of human social behaviour (such as social engagement or social reward, not included in this study) may reveal different association patterns. Fourth, in contrast to preschool scores, analyses of comparable school-age musicality measures revealed effect attenuation. This is consistent with lower study power due to ceiling effects, attrition and/or an increase of non-genetic influences due to a shared school environment. The GWAS of rhythmicity that we investigated in this work is, to date, the largest genetic study of a musicality phenotype. There are ongoing GWAS efforts within the Musicality Genomics Consortium (https://www.mcg.uva.nl/musicgens/), studying other musicality phenotypes, that will become available during the next years. Future research should consider the longitudinal collection of a wide range of musicality and communication phenotypes in powerful multi-ancestry samples to refine and characterise the biological processes underlying the reported findings.

In conclusion, we show that preschool musicality and school-age speech-and syntax-related communication abilities (i) can be predicted by polygenic load for rhythmicity and that, within a developmental context, preschool musicality and speech-and syntax-related communication are linked through shared genetic influences that are (ii) partially attributable to rhythmicity and (iii) independent of cognition and working memory. Consequently, preschool musical engagement involving rhythmic skills may represent a genetic precursor of school-age speech-related communication skills above and beyond general cognition. Our findings strengthen the support for music-related preschool intervention programmes for children with intelligibility-, fluency-and syntax-related communication problems.

## Data availability

The data used are available through a fully searchable data dictionary (http://www.bristol.ac.uk/alspac/researchers/our-data/). Access to ALSPAC data can be obtained as described within the ALSPAC data access policy (http://www.bristol.ac.uk/alspac/researchers/access/).

## Code availability

This study used openly available software. Specifically, PLINK (PLINK v1.9, https://www.cog-genomics.org/plink/1.9/), PRScs (https://github.com/getian107/PRScs). Analyses were mainly performed in R (version 4.1.1) using the following R packages: lavaan (v0.6-14), genomicSEM (v0.0.5), rcompanion (v2.4.30), dplyr (v1.0.8), nFactors (v2.4.1), Matrix (v1.4-1). Requests for scripts or other analysis details can be sent via email to the corresponding authors

## Acknowledgements

The UK Medical Research Council and Wellcome (Grant ref: 217065/Z/19/Z) and the University of Bristol provide core support for ALSPAC. This publication is the work of the authors and they will serve as guarantors for the contents of this paper. A comprehensive list of grant funding is available on the ALSPAC website: http://www.bristol.ac.uk/alspac/external/documents/grant-acknowledgements.pdf. L.D.H., E.V., S.E.F. and B.S.P. were fully supported by the Max Planck Society. B.S.P. is also supported by the European Commission (HORIZON-HLTH-2021 R2D2-MH, 101057385). M.L. received grant support from the National Institute of Mental Health and the National Center for Complementary and Integrative Health under award number R33MH123029. R.L.G received grant support from the National Institute on Deafness and Other Communication Disorders of the National Institutes of Health (NIH) under award number R01DC016977. The content is solely the responsibility of the authors and does not necessarily represent the official views of the NIH. The authors are extremely grateful to all the families who took part in this study, the midwives for their help in recruiting them, and the whole ALSPAC team, which includes interviewers, computer and laboratory technicians, clerical workers, research scientists, volunteers, managers, receptionists and nurses. We thank Chin Yang Shapland for her helpful comments.

## Author contributions

LDH performed the main analysis and BSP supervised the research. BSP. developed the study concept and, LDH contributed to the study design. LDH, EV, JRV and CF analysed the genetic and phenotypic data. LDH and BSP wrote the manuscript. LDH, EV, AO, JRV, CF, ML, SEF, RLG and BSP read and commented on the manuscript.

## References

1. Levinson, S. C. Interactional Foundations of Language: The Interaction Engine Hypothesis. in Human Language (ed. Hagoort, P.) 189–200 (The MIT Press, 2019). doi:10.7551/mitpress/10841.003.0018.

2. ASHA Practice Policy. American Speech-Language-Hearing Association https://www.asha.org/policy.

3. Social Communication. American Speech-Language-Hearing Association https://www.asha.org/public/speech/development/social-communication/.

4. Dall, M., Fellinger, J. & Holzinger, D. The link between social communication and mental health from childhood to young adulthood: A systematic review. Frontiers in Psychiatry 13, (2022).

5. Bishop, D. V. M. Genes, Cognition, and Communication. Annals of the New York Academy of Sciences 1156, 1–18 (2009).

6. Hwa-Froelich, D. A. Social Communication Development and Disorders. (Psychology Press, 2014).

7. Winkler, I., Háden, G. P., Ladinig, O., Sziller, I. & Honing, H. Newborn infants detect the beat in music. Proceedings of the National Academy of Sciences 106, 2468–2471 (2009).

8. Honing, H. On the biological basis of musicality. Annals of the New York Academy of Sciences 1423, 51–56 (2018).

9. Gingras, B., Honing, H., Peretz, I., Trainor, L. J. & Fisher, S. E. Defining the biological bases of individual differences in musicality. Philosophical Transactions of the Royal Society B: Biological Sciences 370, 20140092 (2015).

10. Oesch, N. Music and Language in Social Interaction: Synchrony, Antiphony, and Functional Origins. Frontiers in Psychology 10, (2019).

11. Cirelli, L. K., Trehub, S. E. & Trainor, L. J. Rhythm and melody as social signals for infants. Annals of the New York Academy of Sciences 1423, 66–72 (2018).

12. Lense, M. D., Shultz, S., Astésano, C. & Jones, W. Music of infant-directed singing entrains infants’ social visual behavior. Proceedings of the National Academy of Sciences 119, e2116967119 (2022).

13. Moritz, C., Yampolsky, S., Papadelis, G., Thomson, J. & Wolf, M. Links between early rhythm skills, musical training, and phonological awareness. Read Writ 26, 739–769 (2013).

14. Nayak, S. et al. The Musical Abilities, Pleiotropy, Language, and Environment (MAPLE) Framework for Understanding Musicality-Language Links Across the Lifespan. Neurobiology of Language 3, 615–664 (2022).

15. Swaminathan, S. & Schellenberg, E. G. Musical ability, music training, and language ability in childhood. *Journal of Experimental Psychology: Learning*, Memory, and Cognition 46, 2340 (2019).

16. Bishop, D. V. M., Laws, G., Adams, C. & Norbury, C. F. High Heritability of Speech and Language Impairments in 6-year-old Twins Demonstrated Using Parent and Teacher Report. Behav Genet 36, 173–184 (2006).

17. Wesseldijk, L. W., Ullén, F. & Mosing, M. A. Music and Genetics. Neuroscience & Biobehavioral Reviews 152, 105302 (2023).

18. Niarchou, M. et al. Genome-wide association study of musical beat synchronization demonstrates high polygenicity. Nat Hum Behav 6, 1292–1309 (2022).

19. Alagöz, G., et al. The shared genetic architecture and evolution of human language and musical rhythm. http://biorxiv.org/lookup/doi/10.1101/2023.11.01.564908 (2023) doi:10.1101/2023.11.01.564908.

20. Nayak, S. et al. Musical rhythm abilities and risk for developmental speech-language problems and disorders: epidemiological and polygenic associations. (2024) doi:10.31234/osf.io/kcgp5.

21. Wesseldijk, L. W., Lu, Y., Karlsson, R., Ullén, F. & Mosing, M. A. A comprehensive investigation into the genetic relationship between music engagement and mental health. Transl Psychiatry 13, 1–8 (2023).

22. Gustavson, D. E. et al. Exploring the genetics of rhythmic perception and musical engagement in the Vanderbilt Online Musicality Study. Annals of the New York Academy of Sciences 1521, 140–154 (2023).

23. Wesseldijk, L. W., Abdellaoui, A., Gordon, R. L., Ullén, F. & Mosing, M. A. Using a polygenic score in a family design to understand genetic influences on musicality. Sci Rep 12, 14658 (2022).

24. Marees, A. T. et al. Genetic correlates of socio-economic status influence the pattern of shared heritability across mental health traits. Nat Hum Behav 5, 1065– 1073 (2021).

25. Ladányi, E., Persici, V., Fiveash, A., Tillmann, B. & Gordon, R. L. Is atypical rhythm a risk factor for developmental speech and language disorders? WIREs Cognitive Science 11, e1528 (2020).

26. 26. Fraser, A., et al. Cohort Profile: The Avon Longitudinal Study of Parents and Children: ALSPAC mothers cohort. International Journal of Epidemiology 42, 97– 110 (2013).

27. Boyd, A. et al. Cohort Profile: The ‘Children of the 90s’—the index offspring of the Avon Longitudinal Study of Parents and Children. International Journal of Epidemiology 42, 111–127 (2013).

28. Castles, A., Rastle, K. & Nation, K. Ending the Reading Wars: Reading Acquisition From Novice to Expert. Psychol Sci Public Interest 19, 5–51 (2018).

29. CDC. What developmental milestones is your 1-year-old reaching? Centers for Disease Control and Prevention https://www.cdc.gov/ncbddd/actearly/milestones/milestones-1yr.html (2022).

30. Skuse, D. H., Mandy, W. P. L. & Scourfield, J. Measuring autistic traits: heritability, reliability and validity of the Social and Communication Disorders Checklist. The British Journal of Psychiatry 187, 568–572 (2005).

31. Bishop, D. V. M. Development of the Children’s Communication Checklist (CCC): A Method for Assessing Qualitative Aspects of Communicative Impairment in Children. Journal of Child Psychology and Psychiatry 39, 879–891 (1998).

32. Wechsler, D., Golombok, S. & Rust, J. WISC-III UK Wechsler intelligence scale for children: UK manual. Sidcup, UK: The Psychological Corporation (1992).

33. Gathercole, S. E., Willis, C. S., Baddeley, A. D. & Emslie, H. The children’s test of nonword repetition: A test of phonological working memory. Memory 2, 103–127 (1994).

34. Chang, C. C. et al. Second-generation PLINK: rising to the challenge of larger and richer datasets. GigaScience 4, s13742-015-0047–8 (2015).

35. Wray, N. R. et al. From Basic Science to Clinical Application of Polygenic Risk Scores: A Primer. JAMA Psychiatry 78, 101 (2021).

36. Ge, T., Chen, C.-Y., Ni, Y., Feng, Y.-C. A. & Smoller, J. W. Polygenic prediction via Bayesian regression and continuous shrinkage priors. Nat Commun 10, 1776 (2019).

37. Li, J. & Ji, L. Adjusting multiple testing in multilocus analyses using the eigenvalues of a correlation matrix. Heredity 95, 221–227 (2005).

38. Rosseel, Y. lavaan: An R Package for Structural Equation Modeling. J. Stat. Soft. 48, 1–36 (2012).

39. Baron, R. M. & Kenny, D. A. The moderator–mediator variable distinction in social psychological research: Conceptual, strategic, and statistical considerations. Journal of Personality and Social Psychology 51, 1173–1182 (1986).

40. Okbay, A. et al. Polygenic prediction of educational attainment within and between families from genome-wide association analyses in 3 million individuals. Nat Genet 54, 437–449 (2022).

41. Demange, P. A. et al. Investigating the genetic architecture of noncognitive skills using GWAS-by-subtraction. Nat Genet 53, 35–44 (2021).

42. Grotzinger, A. D. et al. Genomic structural equation modelling provides insights into the multivariate genetic architecture of complex traits. Nat Hum Behav 3, 513– 525 (2019).

43. Abdellaoui, A. & Verweij, K. J. H. Dissecting polygenic signals from genome-wide association studies on human behaviour. Nat Hum Behav 5, 686–694 (2021).

44. Polderman, T. J. C. et al. Meta-analysis of the heritability of human traits based on fifty years of twin studies. Nat Genet 47, 702–709 (2015).

45. Müllensiefen, D., Gingras, B., Musil, J. & Stewart, L. The Musicality of Non-Musicians: An Index for Assessing Musical Sophistication in the General Population. PLoS One 9, e89642 (2014).

46. Politimou, N., Stewart, L., Müllensiefen, D. & Franco, F. Music@Home: A novel instrument to assess the home musical environment in the early years. PLOS ONE 13, e0193819 (2018).

47. Lense, M. D., Ladányi, E., Rabinowitch, T.-C., Trainor, L. & Gordon, R. Rhythm and timing as vulnerabilities in neurodevelopmental disorders. Philos Trans R Soc Lond B Biol Sci 376, 20200327 (2021).

48. Cheverud, J. M. A Comparison of Genetic and Phenotypic Correlations. Evolution 42, 958–968 (1988).

49. Linnavalli, T., Putkinen, V., Lipsanen, J., Huotilainen, M. & Tervaniemi, M. Music playschool enhances children’s linguistic skills. Sci Rep 8, 8767 (2018).

50. Moreno, S. et al. Short-Term Music Training Enhances Verbal Intelligence and Executive Function. Psychol Sci 22, 1425–1433 (2011).

