## Supplementary Material for "Preschool musicality is associated with school-age communication abilities through genes related to rhythmicity"

### Index

**Supplementary Note 1.** Description of ALSPAC cohort

Pregnant women resident in Avon, UK with expected dates of delivery between 1st April 1991 and 31st December 1992 were invited to take part in the study[1,2]. The initial number of pregnancies enrolled was 14,541. Of the initial pregnancies, there was a total of 14,676 foetuses, resulting in 14,062 live births and 13,988 children who were alive at 1 year of age. When the oldest children were approximately 7 years of age, an attempt was made to bolster the initial sample with eligible cases who had failed to join the study originally. As a result, when considering variables collected from the age of seven onwards (and potentially abstracted from obstetric notes) there are data available for more than the 14,541 pregnancies mentioned above. The number of new pregnancies not in the initial sample (known as Phase I enrolment) that are currently represented in the released data and reflecting enrolment status at the age of 24 is 906, resulting in an additional 913 children being enrolled (456, 262 and 195 recruited during Phases II, III and IV respectively). The phases of enrolment are described in more detail in the cohort profile paper and its update. The total sample size for analyses using any data collected after the age of seven is therefore 15,447 pregnancies, resulting in 15,658 foetuses. Of these 14,901 children were alive at 1 year of age. Please note that the study website contains details of all the data that is available through a fully searchable data dictionary and variable search tool: http://www.bristol.ac.uk/alspac/researchers/our-data. Ethical approval for the study was obtained from the ALSPAC Ethics and Law Committee and the Local Research Ethics Committees. Consent for biological samples has been collected in accordance with the Human Tissue Act (2004). Informed consent for the use of data collected via questionnaires and clinics was obtained from participants following recommendations of the ALSPAC Ethics and Law Committee at the time.

**Supplementary Note 2.** Verbal cognition-related measures in ALSPAC

Verbal cognition-related measures were extracted from the Focus@8 questionnaire in ALSPAC. Questionnaires were administered to ALSPAC children by trained assessors in 20-minute sessions. These variables were included to capture variance in communication variables over and above the effects of general cognition.

*Verbal abilities at age 9.* Verbal cognition (verbal IQ) measures children's verbal ability (including working memory and verbal comprehension) and was assessed with an abbreviated form of the Wechsler Intelligence Scale for Children (WISC-III)[3] at 9 years (https://closer.ac.uk/cross-study-data-guides/cognitive-measures-guide/alspac-cognition/alspac-age-8-5-wechsler-intelligence-scale-for/). ALSPAC participants completed an abbreviated form of the WISC, which included alternate items from each of the five verbal subtests: i) information (assessing child's factual knowledge), ii) similarities (where similarities between things must be explained, e.g. *in what way are red and blue alike?*), iii) arithmetic (mental arithmetic questions; assessing child's numerical reasoning), iv) vocabulary (child's understanding of the meaning of different words), and v) comprehension (where the child is asked questions about different situations, e.g. *why are names in the telephone book in alphabetical order?*).

*Non-word repetition at age 9.* Non-word repetition was assessed with an adaptation of the Nonword Repetition Test (NWRT)[4] at 9 years, and captures the children's phonological short-term memory and phonological awareness (https://closer.ac.uk/cross-study-data-guides/cognitive-measures-guide/alspac-cognition/alspac-age-8-5-nonword-repetition/). Children were asked to listen to nonsense words and then repeat each item. These were twelve nonsense words, four each of 3, 4 and 5 syllables and conformed to English rules for sound combinations.

**Supplementary Note 3.** Quality control of genetic data in ALSPAC

We carried out standard quality control procedures at the genetic and individual level in PLINK (v1.07)[5], as previously described[6]. We excluded individuals based on sex mismatch (between reported and genetic sex), SNP missingness (>3%), population stratification (non-European genetic ancestry), or interindividual relatedness (IBD > 5%). Genetic variants were excluded based on individual missingness (>1%), Hardy-Weinberg equilibrium deviations ($p<5x10^{-7}$) or low allele frequency (<1%).

**Supplementary Note 4.** Univariate polygenic score (PGS) analyses in ALSPAC

*PGS calculation*

We conducted PGS analyses in ALSPAC using PRS-CS[7], a Bayesian-based approach that adjusts single-nucleotide polymorphism (SNP) effect sizes for linkage disequilibrium by applying a continuous-shrinkage parameter. Here, we selected the auto-option for a fully Bayesian estimation of the shrinkage parameter phi and used the software's default settings (a=1; b=0.5; Markov Chain Monte Carlo iterations n=1,000; burn-in iterations n=500; Markov chain thinning factor=5). We used the 1000 Genomes European reference panel recommended on the software's GitHub page (https://github.com/getian107/PRScs).

PGS were constructed for unrelated ALSPAC children (genomic relatedness<0.125), based on high-quality imputed HapMap 3 SNPs (INFO>0.8, 95%-posterior genotyping probability>0.9, minor allele frequency > 0.5%). Per-allele posterior effect sizes for SNPs were calculated in PRS-CS and, subsequently, PGS scores were calculated in PLINK (v1.9)[8] and, subsequently, Z-standardised.

*Association analysis*

To test for the association between ALSPAC phenotypes and PGS, we fitted linear regression to continuous traits and binomial regression to binary traits. Regression analyses were corrected for age, sex and the first ten ancestry-informative principal components (applied to correct for subtle population differences[9]).

For each phenotype, we fitted two models: a covariate model: $phenotype \sim sex + age + pc_{1 \ldots 10}$, and a PGS model: $phenotype \sim sex + age + pc_{1 \ldots 10} + PGS$. For continuous traits, we assessed $R^2$ to test for association[10], where

*$R^2$* is the difference of *$R^2$* between the covariate and the PGS models. For binary traits, we assessed *Nagelkerke-$R^2$* as the difference between the covariate and the PGS models, computed using the *nagelkerke()* function of *rcompanion* R package (R::rcompanion, v2.4.30)[11].

**Supplementary Note 5.** Phenotype modelling

To study the phenotypic relationships across traits, we applied a data-driven approach using principal component analysis (PCA), exploratory (EFA) and confirmatory factor analysis (CFA)[12]. EFA and CFA models were fitted with a maximum likelihood estimator using both orthogonal (varimax) and oblique (oblimin) rotation.

First, we estimate the optimal number of factors across the phenotypes by carrying out an eigenvalue decomposition (PCA) of the Pearson phenotypic correlation matrix derived from the full sample. The number of factors was then estimated according to the optimal coordinate criterion[13] (R::nFactors, v2.4.1), which applies a joint Kaiser's rule (eigenvalue > 1)[14] and Cattell's scree test[15]. Second, we randomly split the full sample into two independent halves, matching them based on sex and phenotype missingness patterns using the "slice_sample" function in *dplyr* (R::dplyr, v1.0.8)[16]. Third, we fitted an EFA to the first random half of the sample (N=3,048). To approximate the EFA factor structure, we retained standardised EFA factor loadings ($\lambda$), capturing at least 1% of the phenotypic variation ($|\lambda|>0.1$). Fourth, we fitted an CFA to the second random half of the sample (N=3,053) using the structure identified by EFA. We fixed the variance of the latent variables to one. CFA model fit was assessed using the comparative fit index (CFI), the Tucker–Lewis index (TLI), the Root Mean Square Error of Approximation (RMSEA) and the Standardised Root Mean Square Residual (SRMR) parameters. To evaluate the model fit we used the recommended cut-off criteria[17] for a maximum likelihood estimator: CFI and TLI above 0.95, RMSEA below 0.06 and SRMR below 0.08 indicate an optimal fit.

**Supplementary Note 6.** Phenotype transformations

To account for covariate effects in factor analyses within our study, we transformed all scores adjusting for age, sex and the first ten ancestry-informative principal components from the genotyping analysis to correct for population stratification[9]. This was carried out by regressing measures on covariates using logistic (binary traits) or linear (continuous traits) regression, as previously described[12].

For binary traits, we extracted deviance residuals (*resid()* R function) of the logistic regression. For continuous phenotypes, residuals of the linear regression (*resid()* R function) were rank-transformed and regressed on covariates to achieve normality of transformed scores while avoiding a re-introduction of covariate effects[18].

**Supplementary Table 1.** $PGS_{rhythmicity}$ analysis for ALSPAC measures

| Variable | BETA | SE | P | Nagelkerke-R2 | Domain |
|---|---|---|---|---|---|
| Nursery rhymes 0.5Y | 0.097 | 0.026 | $2 \times 10^{-4}$ | 0.003 | Preschool nursery rhymes |
| Nursery rhymes 1.5Y | -0.004 | 0.037 | $9 \times 10^{-1}$ | <0.001 | Preschool nursery rhymes |
| Can sing at least 3 songs 5Y | 0.151 | 0.108 | $2 \times 10^{-1}$ | 0.002 | Preschool musicality |
| Can hum a tune 5Y | 0.231 | 0.067 | $6 \times 10^{-4}$ | 0.007 | Preschool musicality |
| Can clap to a beat 5Y | 0.223 | 0.053 | $2 \times 10^{-5}$ | 0.008 | Preschool musicality |
| Can sing at least 3 songs 6Y | 0.066 | 0.156 | $7 \times 10^{-1}$ | <0.001 | School-age musicality |
| Can hum a tune 6Y | 0.075 | 0.097 | $4 \times 10^{-1}$ | 0.001 | School-age musicality |
| Can clap to a beat 6Y | 0.191 | 0.086 | $3 \times 10^{-2}$ | 0.004 | School-age musicality |
| Can sing at least 3 songs 7Y | 0.286 | 0.159 | $7 \times 10^{-2}$ | 0.007 | School-age musicality |
| Can hum a tune 7Y | 0.324 | 0.141 | $2 \times 10^{-2}$ | 0.010 | School-age musicality |
| Can clap to a beat 7Y | 0.185 | 0.122 | $1 \times 10^{-1}$ | 0.003 | School-age musicality |

| Variable | BETA | SE | P | R2 | Domain |
|---|---|---|---|---|---|
| Intelligibility and fluency 10Y | 0.119 | 0.025 | $1 \times 10^{-6}$ | 0.004 | School-age communication (CCC) |
| Syntax 10Y | 0.022 | 0.008 | $3 \times 10^{-3}$ | 0.002 | School-age communication (CCC) |
| Appropriate initiation 10Y | -0.068 | 0.032 | $3 \times 10^{-2}$ | 0.001 | School-age communication (CCC) |
| Coherence 10Y | 0.050 | 0.026 | $6 \times 10^{-2}$ | 0.001 | School-age communication (CCC) |
| Non-stereotyped conversation 10Y | -0.003 | 0.033 | $9 \times 10^{-1}$ | <0.001 | School-age communication (CCC) |
| Use of conversational context 10Y | 0.021 | 0.028 | $4 \times 10^{-1}$ | <0.001 | School-age communication (CCC) |
| Conversational rapport 10Y | 0.067 | 0.026 | $9 \times 10^{-3}$ | 0.001 | School-age communication (CCC) |
| Pragmatic score 10Y | 0.069 | 0.103 | $5 \times 10^{-1}$ | <0.001 | School-age communication (CCC) |

**Table S1** (continued).

| Variable | BETA | SE | P | R2 | Domain |
|---|---|---|---|---|---|
| SCDC score 8Y | -0.093 | 0.050 | 6 $x10^{-2}$ | 0.001 | School-age social-communication (SCDC) |
| SCDC score 11Y | -0.006 | 0.048 | 9 $x10^{-1}$ | <0.001 | School-age social-communication (SCDC) |
| SCDC score 14Y | -0.078 | 0.050 | 1 $x10^{-1}$ | <0.001 | School-age social-communication (SCDC) |
| SCDC score 17Y | 0.016 | 0.058 | 8 $x10^{-1}$ | <0.001 | School-age social-communication (SCDC) |
| Nonword repetition 9Y (NWRT) | 0.244 | 0.034 | 5 $x10^{-13}$ | 0.010 | School-age cognition-related |
| Verbal IQ 9Y (WISC-III) | -0.545 | 0.224 | 1 $x10^{-2}$ | 0.001 | School-age cognition-related |

**Supplementary Table 2.** Relationships between factors and association analysis of $PGS_{rhythmicity}$ with factors

| Relationships between factors | estimate | se | P |
|---|---|---|---|
| F1 → $PGS_{rhythmicity}$ (a) | 0.096 | 0.021 | $5.6 \times 10^{-6}$ |
| $PGS_{rhythmicity}$ → F2 ($b_{F2}$) | 0.016 | 0.022 | $4.8 \times 10^{-1}$ |
| F1 → F2 ($c_{F1-F2}$) | 0.21 | 0.033 | $5.2 \times 10^{-11}$ |
| $PGS_{rhythmicity}$ → F3 ($b_{F3}$) | 0.068 | 0.019 | $2.0 \times 10^{-4}$ |
| F1 → F3 ($c_{F1-F3}$) | 0.26 | 0.047 | $4.2 \times 10^{-9}$ |
| **Association between F1 and F2 through $PGS_{rhythmicity}$** | **estimate** | **se** | **P** |
| Shared effect (a x $b_{F2}$) | 0.0015 | 0.0022 | $4.9 \times 10^{-1}$ |
| Total effect (a x $b_{F2}$ + $c_{F1-F2}$) | 0.21 | 0.033 | $2.4 \times 10^{-11}$ |
| **Association between F1 and F3 through $PGS_{rhythmicity}$** | **estimate** | **se** | **P** |
| Shared effect (a x $b_{F3}$) | 0.0065 | 0.0021 | $1.6 \times 10^{-3}$ |
| Total effect (a x $b_{F3}$ + $c_{F1-F3}$) | 0.27 | 0.047 | $1.6 \times 10^{-9}$ |

**Supplementary Table 3.** $PGS_{rhythmicity-EA}$ analysis for ALSPAC measures

| Variable | BETA | SE | P | Nagelkerke-R2 | Domain |
|---|---|---|---|---|---|
| Nursery rhymes 0.5Y | 0.094 | 0.026 | $3x10^{-4}$ | 0.003 | Preschool nursery rhymes |
| Nursery rhymes 1.5Y | -0.023 | 0.037 | $5 x10^{-1}$ | <0.001 | Preschool nursery rhymes |
| Can sing at least 3 songs 5Y | 0.146 | 0.108 | $8 x10^{-4}$ | 0.002 | Preschool musicality |
| Can hum a tune 5Y | 0.223 | 0.067 | $1 x10^{-4}$ | 0.007 | Preschool musicality |
| Can clap to a beat 5Y | 0.200 | 0.052 | $3 x10^{-1}$ | 0.006 | Preschool musicality |
| Can sing at least 3 songs 6Y | 0.147 | 0.156 | $5 x10^{-1}$ | 0.002 | School-age musicality |
| Can hum a tune 6Y | 0.061 | 0.097 | $2 x10^{-2}$ | <0.001 | School-age musicality |
| Can clap to a beat 6Y | 0.204 | 0.086 | $9 x10^{-2}$ | 0.005 | School-age musicality |
| Can sing at least 3 songs 7Y | 0.271 | 0.159 | $2 x10^{-2}$ | 0.007 | School-age musicality |
| Can hum a tune 7Y | 0.338 | 0.141 | $4 x10^{-2}$ | 0.011 | School-age musicality |
| Can clap to a beat 7Y | 0.249 | 0.123 | $8 x10^{-4}$ | 0.006 | School-age musicality |

| Variable | BETA | SE | P | R2 | Domain |
|---|---|---|---|---|---|
| Intelligibility and fluency 10Y | 0.110 | 0.025 | $1 x10^{-2}$ | 0.003 | School-age communication (CCC) |
| Syntax 10Y | 0.020 | 0.008 | $2 x10^{-1}$ | 0.001 | School-age communication (CCC) |
| Appropriate initiation 10Y | -0.044 | 0.032 | $1 x10^{-1}$ | <0.001 | School-age communication (CCC) |
| Coherence 10Y | 0.040 | 0.026 | $8 x10^{-1}$ | <0.001 | School-age communication (CCC) |
| Non-stereotyped conversation 10Y | 0.009 | 0.033 | $4 x10^{-1}$ | <0.001 | School-age communication (CCC) |
| Use of conversational context 10Y | 0.026 | 0.028 | $5 x10^{-2}$ | <0.001 | School-age communication (CCC) |
| Conversational rapport 10Y | 0.050 | 0.026 | $4 x10^{-1}$ | 0.001 | School-age communication (CCC) |
| Pragmatic score 10Y | 0.079 | 0.103 | $4 x10^{-2}$ | <0.001 | School-age communication (CCC) |

**Table S3** continued.

| Variable | BETA | SE | P | R2 | Domain |
|---|---|---|---|---|---|
| SCDC score 8Y | -0.100 | 0.050 | $9 \times 10^{-1}$ | 0.001 | School-age social-communication (SCDC) |
| SCDC score 11Y | -0.006 | 0.048 | $7 \times 10^{-2}$ | <0.001 | School-age social-communication (SCDC) |
| SCDC score 14Y | -0.091 | 0.050 | $1 \times 10^{-0}$ | 0.001 | School-age social-communication (SCDC) |
| SCDC score 17Y | -0.001 | 0.058 | $7 \times 10^{-4}$ | <0.001 | School-age social-communication (SCDC) |
| Nonword repetition 9Y (NWRT) | -0.089 | 0.026 | $7 \times 10^{-14}$ | 0.002 | School-age cognition-related |
| Verbal IQ 9Y (WISC-III) | 0.253 | 0.034 | $4 \times 10^{-2}$ | 0.010 | School-age cognition-related |

**Supplementary Table 4.** Relationships between factors and association analysis of $PGS_{rhythmicity\text{-}EA}$ with factors

| Relationships between factors | estimate | se | P |
|---|---|---|---|
| F1 → $PGS_{rhythmicity\text{-}EA}$ (a) | 0.087 | 0.021 | $4.3 \times 10^{-5}$ |
| $PGS_{rhythmicity\text{-}EA}$ → F2 ($b_{F2}$) | 0.027 | 0.022 | $2.1 \times 10^{-1}$ |
| F1 → F2 ($c_{F1\text{-}F2}$) | 0.21 | 0.033 | $7.1 \times 10^{-11}$ |
| $PGS_{rhythmicity\text{-}EA}$ → F3 ($b_{F3}$) | 0.061 | 0.019 | $7.5 \times 10^{-4}$ |
| F1 → F3 ($c_{F1\text{-}F3}$) | 0.27 | 0.047 | $3.1 \times 10^{-9}$ |
| **Association between F1 and F2 through $PGS_{rhythmicity\text{-}EA}$** | **estimate** | **se** | **P** |
| Shared effect (a x $b_{F2}$) | 0.0024 | 0.0020 | $2.3 \times 10^{-1}$ |
| Total effect (a x $b_{F2}$ + $c_{F1\text{-}F2}$) | 0.21 | 0.033 | $2.9 \times 10^{-11}$ |
| **Association between F1 and F3 through $PGS_{rhythmicity\text{-}EA}$** | **estimate** | **se** | **P** |
| Shared effect (a x $b_{F3}$) | 0.0052 | 0.0019 | $4.7 \times 10^{-3}$ |
| Total effect (a x $b_{F3}$ + $c_{F1\text{-}F3}$) | 0.27 | 0.047 | $1.5 \times 10^{-9}$ |

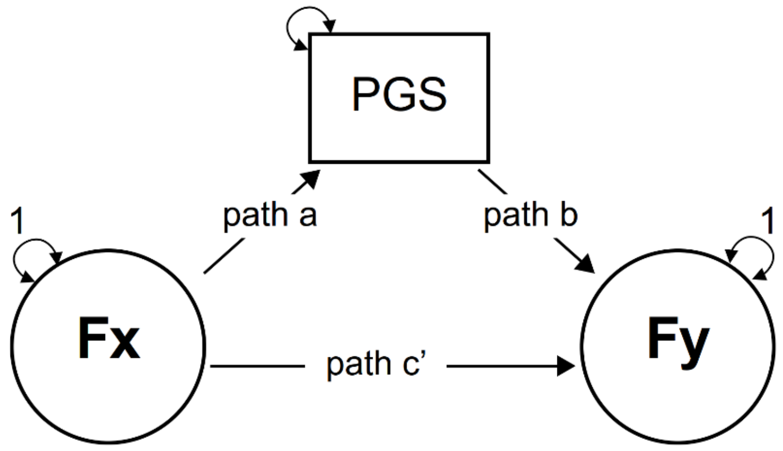

**Supplementary Figure 1.** Mediation methodology applied to factor structures

To test for the association of polygenic scores (PGS) and their relevance in explaining the relationships between factors, we applied a methodology analogous to mediation analysis[19] embedded within structural equation modelling. To do so, we regress (path c’) the outcome phenotypic factor (Fy) against the predicted phenotypic factor (Fx). Simultaneously, the outcome phenotypic factor (Fy) is regressed (path b) against PGS and, in turn, PGS is regressed (path a) against the predictive phenotypic factor (Fx). We computed the shared effect (a*b), captured by the indirect effect (hereforth referred to as shared effect with PGS) within a mediation framework, and the total effect (a*b + c’). Paths were computed in lavaan and SEs were calculated using bootstrapping, following guidelines (https://lavaan.ugent.be/tutorial/mediation.html).

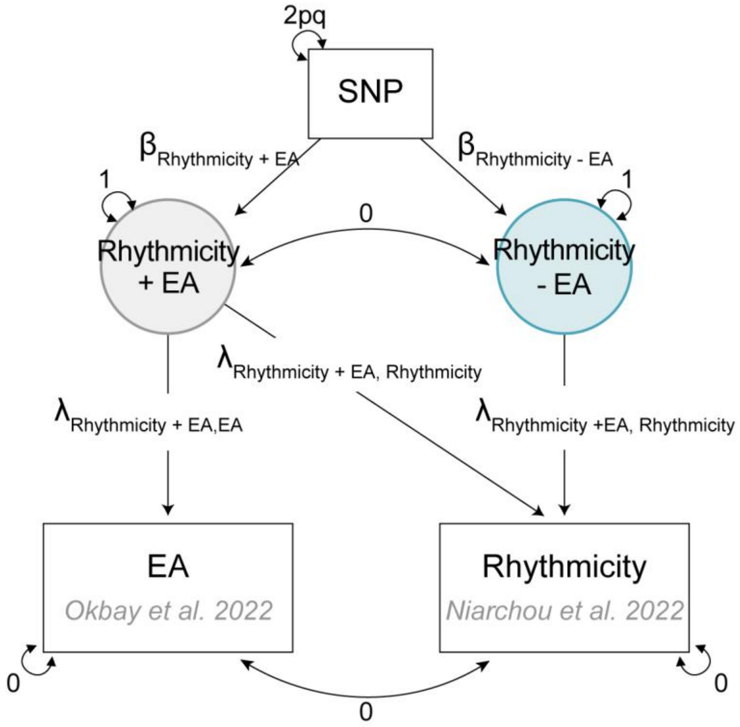

**Supplementary Figure 2.** GWAS-by-subtraction model

This analysis used the methodology described by Demange and colleagues [20]. Circles (*Rhythmicity + EA* and *Rhythmicity – EA*) represent latent (unobserved) variables. Squares (SNP, EA, Rhythmicity) represent observed variables based on GWAS summary statistics. Rhythmicity GWAS summary statistics were extracted from Niarchou et al. 2022[21] and EA GWAS summary statistics were obtained from EA4[22] removing individuals from 23andMe and ALSPAC mothers (to avoid sample overlap). In the model, the variances of the two traits (rhythmicity and EA) are fixed to zero to ensure all the variance is explained by the two latent variables. Abbreviations: EA (educational attainment), SNP (single nucleotide polymorphism).

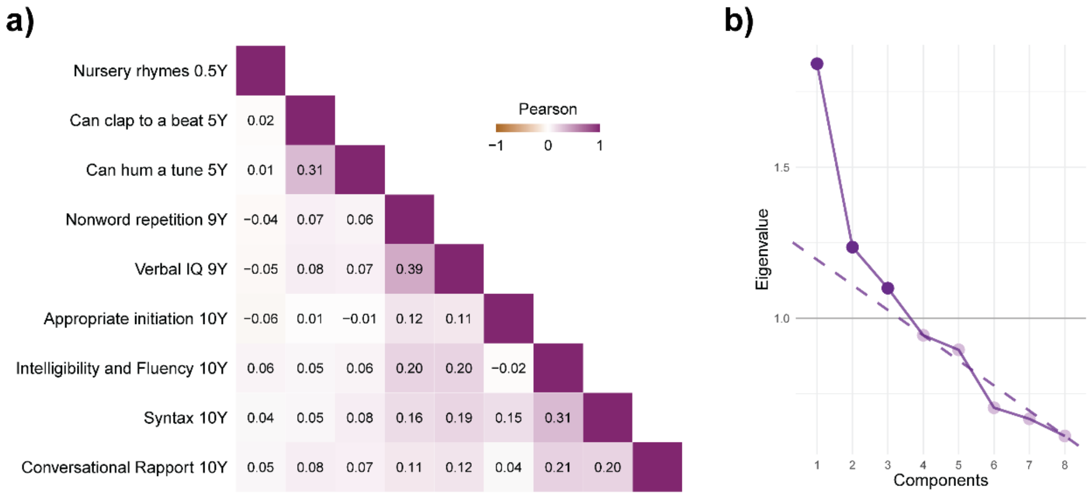

**Supplementary Figure 3.** Eigenvalue decomposition of the phenotypic correlation matrix across $\text{PGS}_{\text{rhythmicity}}$-associated phenotypes

a) Phenotypic correlation matrix. b) Scree plot of eigenvalue decomposition of the phenotypic correlation matrix. The dashed purple line represents the scree, calculated using the optimal coordinate criterion from the nFactors R package[13].

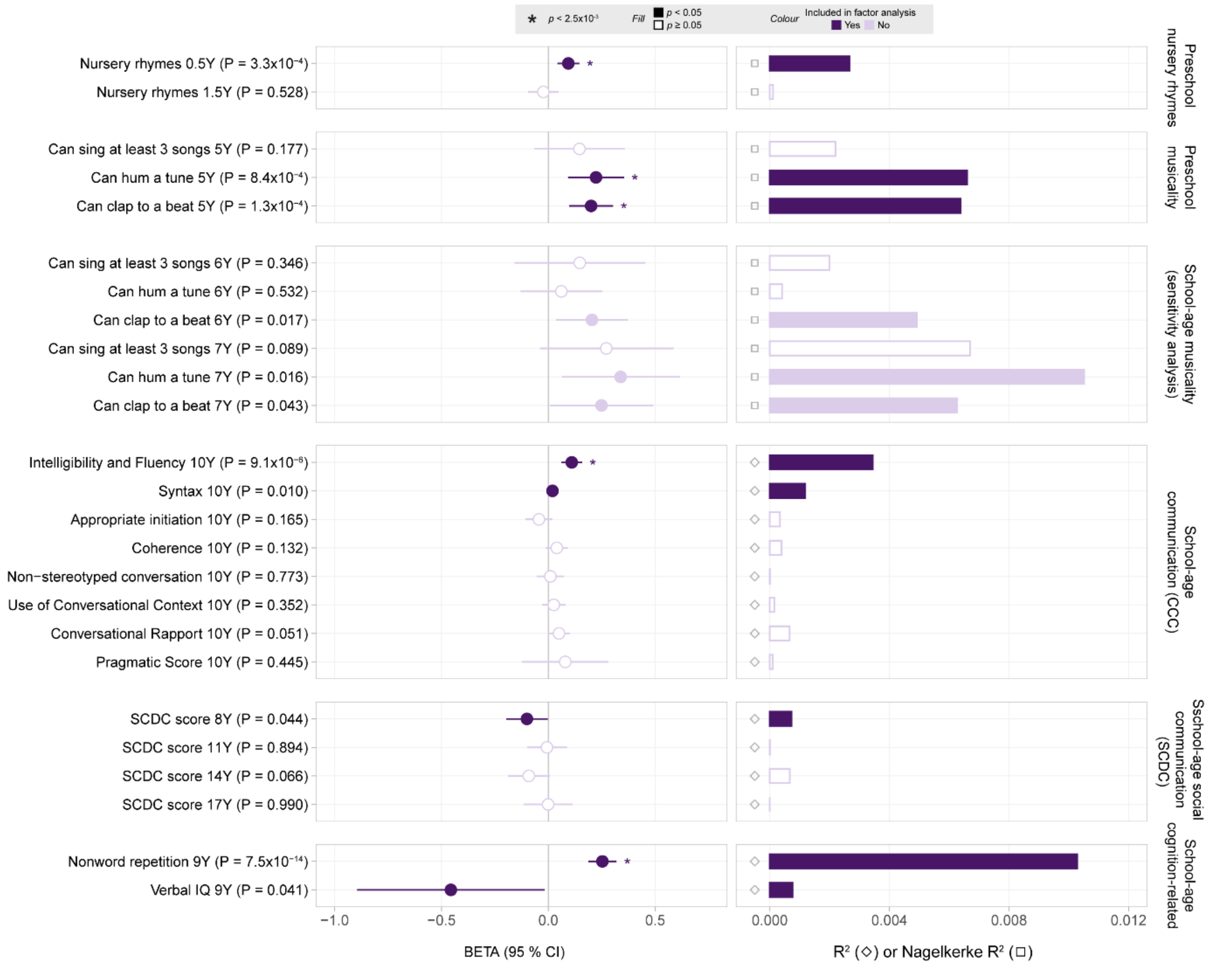

**Supplementary Figure 4.** PGS association analysis of $PGS_{rhythmicity\text{-}EA}$ with ALSPAC phenotypes

Beta estimates are shown as circles with their corresponding 95% confidence intervals. Variance explained for each phenotype is shown as bars and expressed as the regression $R^2$ for continuous traits (represented by an empty grey diamond) and, in analogy, by *Nagelkerke*-$R^2$ for binary traits (represented by an empty grey square). Filled circles/bars and empty circles/bars represent phenotypes with an association with $PGS_{rhythmicity}$ of $p<0.05$ and $p \geq 0.05$, respectively. Estimates are shown in dark purple if they were included in subsequent factor analysis and in light purple otherwise. If a phenotype passed multiple-testing threshold of $2.5 \times 10^{-3}$ this was indicated with an asterisk in the beta coefficient. Abbreviations: SCDC (Social Communication Difficulties Checklist), CCC (Children’s Communication Checklist).

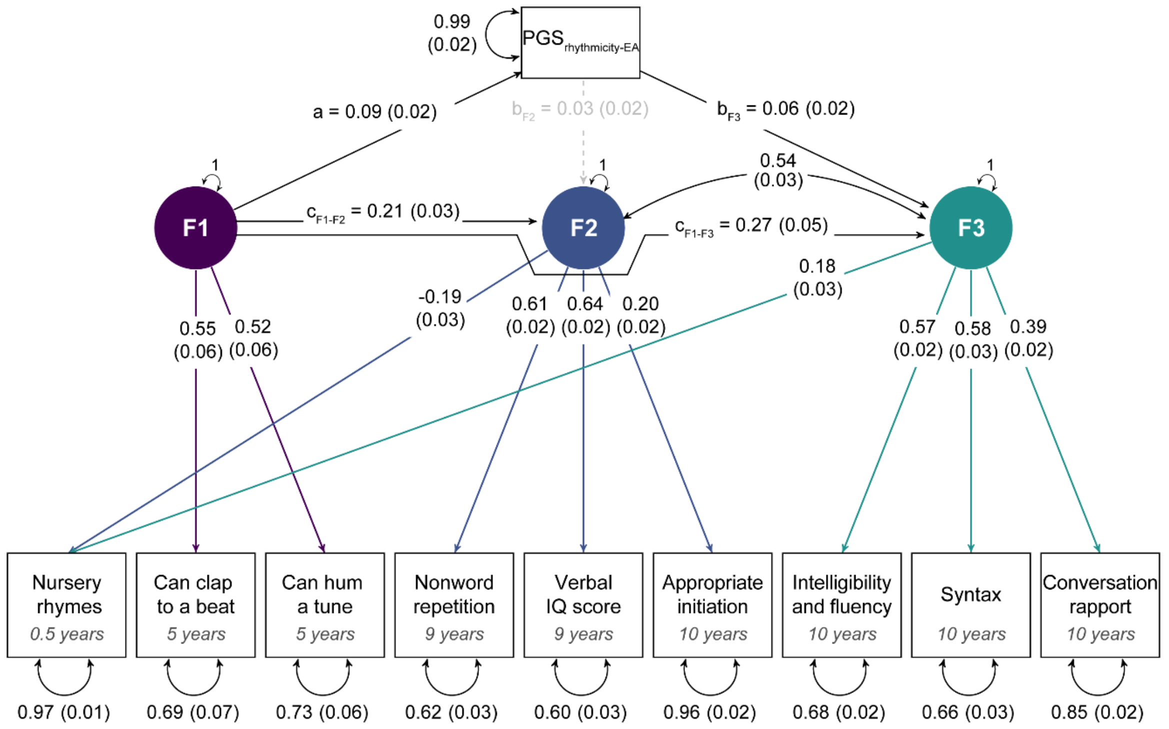

**Supplementary Figure 5.** Genetic characterisation of phenotypic structures with $PGS_{rhythmicity\text{-}EA}$.

Path diagram depicting the best-fitting CFA phenotypic model. Estimates are shown with their corresponding SEs. Observed measures are represented by squares and latent variables by circles. Coloured single-headed arrows define factor loadings with $p \leq 0.05$. Double-headed black arrows represent the variance of each phenotype and factor correlations. Grey dotted and black solid single-headed arrows define relationships between factors and with $PGS_{rhythmicity\text{-}EA}$ with $p > 0.05$ and $p \leq 0.05$, respectively. The shared genetic effect between F1 and F3, as captured by $PGS_{rhythmicity\text{-}EA}$, is estimated as $a^* b_{F3}$ and the total effect between F1 and F3 as $a^* b_{F3} + c_{F1\text{-}F3}$. The shared genetic effect between F1 and F2, as captured by $PGS_{rhythmicity\text{-}EA}$, is estimated as $a^* b_{F2}$ and the total effect as $a^* b_{F2} + c_{F1\text{-}F2}$.

#### Supplementary References

1. Fraser, A. *et al.* Cohort Profile: The Avon Longitudinal Study of Parents and Children: ALSPAC mothers cohort. *International Journal of Epidemiology* **42**, 97–110 (2013).
2. Boyd, A. *et al.* Cohort Profile: The 'Children of the 90s'—the index offspring of the Avon Longitudinal Study of Parents and Children. *International Journal of Epidemiology* **42**, 111–127 (2013).
3. Wechsler, D., Golombok, S. & Rust, J. WISC-III UK Wechsler intelligence scale for children: UK manual. *Sidcup, UK: The Psychological Corporation* (1992).
4. Gathercole, S. E., Willis, C. S., Baddeley, A. D. & Emslie, H. The children's test of nonword repetition: A test of phonological working memory. *Memory* **2**, 103–127 (1994).
5. Purcell, S. *et al.* PLINK: A Tool Set for Whole-Genome Association and Population-Based Linkage Analyses. *The American Journal of Human Genetics* **81**, 559–575 (2007).
6. Verhoef, E. *et al.* Disentangling polygenic associations between attention-deficit/hyperactivity disorder, educational attainment, literacy and language. *Transl Psychiatry* **9**, 1–12 (2019).
7. Ge, T., Chen, C.-Y., Ni, Y., Feng, Y.-C. A. & Smoller, J. W. Polygenic prediction via Bayesian regression and continuous shrinkage priors. *Nat Commun* **10**, 1776 (2019).
8. Chang, C. C. *et al.* Second-generation PLINK: rising to the challenge of larger and richer datasets. *GigaScience* **4**, s13742-015-0047–8 (2015).
9. Price, A. L. *et al.* Principal components analysis corrects for stratification in genome-wide association studies. *Nat Genet* **38**, 904–909 (2006).
10. Choi, S. W., Mak, T. S. H. & O'Reilly, P. F. A guide to performing Polygenic Risk Score analyses. *Nature protocols* **15**, 2759 (2020).
11. Mangiafico, S. S. *rcompanion: Functions to Support Extension Education Program Evaluation*. (Rutgers Cooperative Extension, 2023).
12. de Hoyos, L. *et al.* Structural models of genome-wide covariance identify multiple common dimensions in autism. *Nat Commun* **15**, 1770 (2024).
13. Raîche, G., Walls, T. A., Magis, D., Riopel, M. & Blais, J.-G. Non-graphical solutions for Cattell's scree test. *Methodology: European Journal of Research Methods for the Behavioral and Social Sciences* **9**, 23 (2013).
14. Kaiser, H. F. The Application of Electronic Computers to Factor Analysis. *Educational and Psychological Measurement* **20**, 141–151 (1960).

15. Cattell, R. B. The Scree Test For The Number Of Factors. *Multivariate Behavioral Research* **1**, 245–276 (1966).
16. Wickham, H., François, R., Henry, L. & Müller, K. *dplyr: A Grammar of Data Manipulation*. (2022).
17. Hu, L. & Bentler, P. M. Cutoff criteria for fit indexes in covariance structure analysis: Conventional criteria versus new alternatives. *Structural Equation Modeling: A Multidisciplinary Journal* **6**, 1–55 (1999).
18. Sofer, T. *et al.* A fully adjusted two-stage procedure for rank-normalization in genetic association studies. *Genetic Epidemiology* **43**, 263–275 (2019).
19. Baron, R. M. & Kenny, D. A. The moderator–mediator variable distinction in social psychological research: Conceptual, strategic, and statistical considerations. *Journal of Personality and Social Psychology* **51**, 1173–1182 (1986).
20. Demange, P. A. *et al.* Investigating the genetic architecture of noncognitive skills using GWAS-by-subtraction. *Nat Genet* **53**, 35–44 (2021).
21. Niarchou, M. *et al.* Genome-wide association study of musical beat synchronization demonstrates high polygenicity. *Nat Hum Behav* **6**, 1292–1309 (2022).
22. Okbay, A. *et al.* Polygenic prediction of educational attainment within and between families from genome-wide association analyses in 3 million individuals. *Nat Genet* **54**, 437–449 (2022).
